# Participatory Systems Mapping and Experimental Games to Explore Biosecurity Adoption in Broiler Production in Bangladesh

**DOI:** 10.64898/2026.03.20.712586

**Authors:** Ibrahim Khalil, Md. Nurul Alam, Sajjad Hossain, Md. Yeasin Arafat, Md. Habibur Rahman, A.K.M. Mostafa Anower

**Affiliations:** Field Disease Investigation Laboratory, Barishal, Department of Livestock Services, Ministry of Fisheries and Livestock, Bangladesh; Faculty of Animal Science and Veterinary Medicine, Sher-e-Bangla Agricultural University, Dhaka, Bangladesh; Department of Microbiology and Public Health, Patuakhali Science and Technology University, Patuakhali, Bangladesh; AMR MPTF Bangladesh Project, World Organisation for Animal Health (WOAH), Dhaka, Bangladesh

**Keywords:** Antimicrobial resistance (AMR), Biosecurity, Participatory systems mapping (CLDs), Experimental games, Broiler production

## Abstract

**Introduction:** Antimicrobial Resistance (AMR) presents a critical public health challenge, particularly in smallholder broiler farming, where antibiotics are often used preventively in the absence of effective biosecurity measures.

**Objective:** This study investigates the adoption of biosecurity practices as a sustainable alternative to antibiotics through Participatory Systems Mapping and Experimental Games.

**Methods:** A participatory mixed-methods study was conducted in southern Bangladesh (September 2024-June 2025). Causal Loop Diagrams (CLDs) were co-created with farmers, dealers, and veterinary officers. Ten broiler farmers from single village were selected via purposive and snowball sampling. Experimental games simulated four production cycles where farmers chose Option A (biosecurity, adopters) or Option B (antibiotics, non-adopters) after several interactive trainings. Key metrics including biosecurity compliance (0-12 scale), mortality, FCR, antibiotic use, outbreak history, and economic outcomes were recorded.

**Results:** CLD analysis revealed a reinforcing loop of increased antibiotic reliance driven by fear of mortality, and balancing loops involving training, biosecurity practices, and consumer incentives to reduce use. Five farmers chose Option A, and both groups remained stable until Round 4. Adopters had flock sizes of 800-2000 birds (non-adopters, 600-1000; mean for both = 1000), were younger, and more educated compared to non-adopters. At baseline, both groups had similar biosecurity scores (0). Adopters had higher mean outbreaks (2 vs. 1.4), mortality (5.6 vs. 4.2), antibiotic use (3.6 vs. 3), and FCR (1.8 vs. 1.6) compared to non-adopters. By Round 4, adopters improved biosecurity scores by 125%, eliminated outbreaks, reduced mortality by 52.6%, stopped antibiotic use, improved FCR by 13.3%, and gained 71.72% profit per bird compared to non-adopters. Non-adopters, influenced by adopters, increased biosecurity scores by 25%, reducing outbreaks, mortality, antibiotic use, and FCR. Adopters also increased direct sales to consumers, yielding a 10%-16% profit gain per bird each round.

**Conclusion:** This study highlights the successful adoption of biosecurity practices by farmers, replacing antibiotics and improving production outcomes. Farmer-driven adoption of these practices fosters long-term sustainability and supports a healthier planet within the One Health framework.

## Introduction

Antimicrobial Resistance (AMR) is a significant global threat to public health, affecting both human and animal health[1-5]. In Bangladesh, the broiler poultry sector highlights this issue, where small-scale farms heavily rely on antibiotics, with 98% of commercial farms using them prophylactically to prevent disease[6-8]. Weak biosecurity practices contribute to frequent outbreaks [9-12], increasing antibiotic dependence and fostering the development of drug-resistant pathogens that threaten animal health, farm workers, and consumers through zoonotic transmission and foodborne exposure [3].

The adoption of biosecurity measures in poultry farming is essential for mitigating AMR, improving poultry health, and enhancing farm profitability[12, 13]. Regular cleaning and disinfection are crucial for eliminating pathogens and preventing disease outbreaks [14]. Physical barriers, such as fences to prevent stray animals and separate footwear for different farm zones, help control disease vectors [15]. A minimum 15-day gap between batches for environmental sanitation further reduces disease risks [16]. Farms that implement these measures report lower mortality rates, with some achieving less than 2% mortality in the first week, compared to higher rates in farms lacking such practices [17]. These measures also lead to a significant reduction in antibiotic use, as seen in broiler farms where only 17.4% of farms employing biosecurity measures used antimicrobials three times, compared to higher usage in non-compliant farms [17]. A scoping review further supports the link between improved biosecurity and reduced antibiotic use in poultry [18]. However, the effectiveness of these measures can vary based on implementation quality, active farmer participation, and continuous monitoring, highlighting the need for a comprehensive farm management approach.

Engaging farmers in the community fosters confidence, accountability, and cooperation, promoting biosecurity compliance, reducing antibiotic use, and improving long-term outcomes[13]. Participatory training approaches are crucial for building this confidence, as they effectively communicate complex concepts like microbial resistance and biosecurity, which traditional methods often fail to do[19, 20]. Interactive sessions create a collaborative learning environment where farmers can actively participate, ask questions, and share experiences, enhancing knowledge retention and practice adoption. Visual aids and hands-on activities, such as using glowing substances to demonstrate contamination risks, make biosecurity measures more tangible and relatable [21]. However, challenges remain in translating knowledge into consistent practice, as factors such as gender, education, and initial attitudes influence training effectiveness [22, 23]. Addressing these barriers is essential for the sustained adoption of biosecurity practices and achieving higher adoption rates in poultry farming.

Rising awareness of antibiotic residues has led consumers in Bangladesh to prefer antibiotic-free meat, driving market demand [23]. Studies show that 26% of poultry samples contain antibiotic traces, posing health risks [24, 25]. This demand provides farmers with an opportunity to adopt biosecurity measures and avoid antibiotic use. Farmers who implement antibiotic-free practices benefit from higher prices and improved market access, particularly through e-commerce platforms [26]. This transition also supports income generation for low-income households, contributing to poverty alleviation. Entrepreneurs are responding by rearing antibiotic-free broilers and using online marketing to reach consumers, benefiting both the economy and public health.

Against this backdrop, the present study aims to: (i) identify the feedback loops between antibiotic use, disease vulnerability, and biosecurity practices through Causal Loop Diagrams (CLDs), (ii) evaluate the impact of biosecurity adoption on production outcomes, including mortality rates, feed conversion ratio (FCR), and profitability, and (iii) examine how direct consumer sales contribute to farm profitability by enhancing market access. By employing these methods, this study investigates biosecurity adoption as a sustainable alternative to antibiotics, aligned with the One Health framework, and emphasizes community engagement and integrated training for ensuring long-term sustainability in poultry farming.

## Materials & Methods

### Study Location

The study was conducted in a village (Kawarchar) in Karnakathi union, in Barishal Sadar sub-district, Barishal District, Bangladesh, located at approximately 22.6553° N latitude and 90.3718° E longitude. Barishal is a low-lying deltaic region interspersed with rivers and canals, part of the Ganges-Brahmaputra-Meghna river system,[27, 28] and is primarily agricultural farming playing a key economic role [29, 30]. Barishal Sadar sub-district covers 256.45 square kilometers, with 147 villages and a population of 237,211 people (117,895 male,119,312 female). There are 520 broiler, 739 layer, and 124 Sonali small and medium-scale poultry farms (Personal communication, Sub-district Veterinary Hospital in Barishal Sadar, 2025)

### Model Poultry Village

In November 2023, the Department of Livestock Services (DLS), Bangladesh, supported by Food and Agriculture Organization-Emergency Centre for Transboundary Animal Diseases (FAO-ECTAD), launched a pilot project funded by Fleming Fund across ten sub-districts to encourage poultry farmers to adopt biosecurity measures and promote antibiotic-free farming under the U2C I & II and BARA Model Poultry Farm Pilot Project. In Barishal, 20 farmers from Barishal Sadar and Banaripara sub-districts were selected for this initiative. Due to the scattered nature of the farms, the Field Disease Investigation Laboratory (FDIL), Barishal, under the DLS, faced challenges in continuous monitoring in Barishal. To address this, FDIL Barishal designated a specific village as the Model Poultry Village, based on the U2C I & II and BARA training modules, encouraging farmers to adopt biosecurity-based poultry farming practices.

### U2C (Upazila to Community)

The Upazila (Sub-district) to Community (U2C) program, launched in 2016 by the Bangladesh Department of Livestock Services (DLS) in collaboration with FAO and USAID, aims to strengthen rural animal health by bridging the gap between Upazila veterinary services and community-level farmers [5, 31]. The program focuses on training government field staff to educate farmers on vaccination, biosecurity, and disease surveillance. Initially piloted in nine Upazilas, the program has expanded to cover 126 and is designed to reach all 496 Upazilas across Bangladesh. Additionally, U2C’s initiatives contribute to the broader One Health approach and antimicrobial resistance (AMR) reduction strategies.

### BARA (Bangladesh AMR Response Alliance)

The Bangladesh AMR Response Alliance (BARA) is a group of health professionals including veterinarians, physicians, researchers, and policymakers, following the One Health approach to develop guidelines [5] and awareness tools for responsible antimicrobial use in humans and animals including aquaculture [32]. BARA raises awareness about AMR risks and advocates for best practices in antibiotic stewardship, infection prevention, and disease surveillance, fostering cross-sector collaboration to protect public health and preserve antimicrobial efficacy.

### Village and Farms selection

In the U2C I & II and BARA Model Poultry Farm Pilot Project in Barishal, we identified a motivated farmer in a village of Barishal Sadar sub-district and arranged a meeting with 20 farmers in the farmer’s yard. During this meeting, we applied participatory disease searching (PDS) tools learned from U2C, including structured and semi-structured interviews, focus group discussions (FGDs), simple and pairwise rankings, disease matrix, proportional piling observation, mapping (triangulation & transect work), timeline, and surveillance. Following this, we selected 10 farmers from the 20 in the Model Poultry Village to participate in the experimental game. These farmers were chosen based on convenience, with priority given to those with flock sizes greater than 500 broilers, younger age, higher education, and a willingness to participate. The remaining 10 farmers contributed to the creation of Causal Loop Diagrams (CLDs) (**Fig 1**).

**Fig 1.**
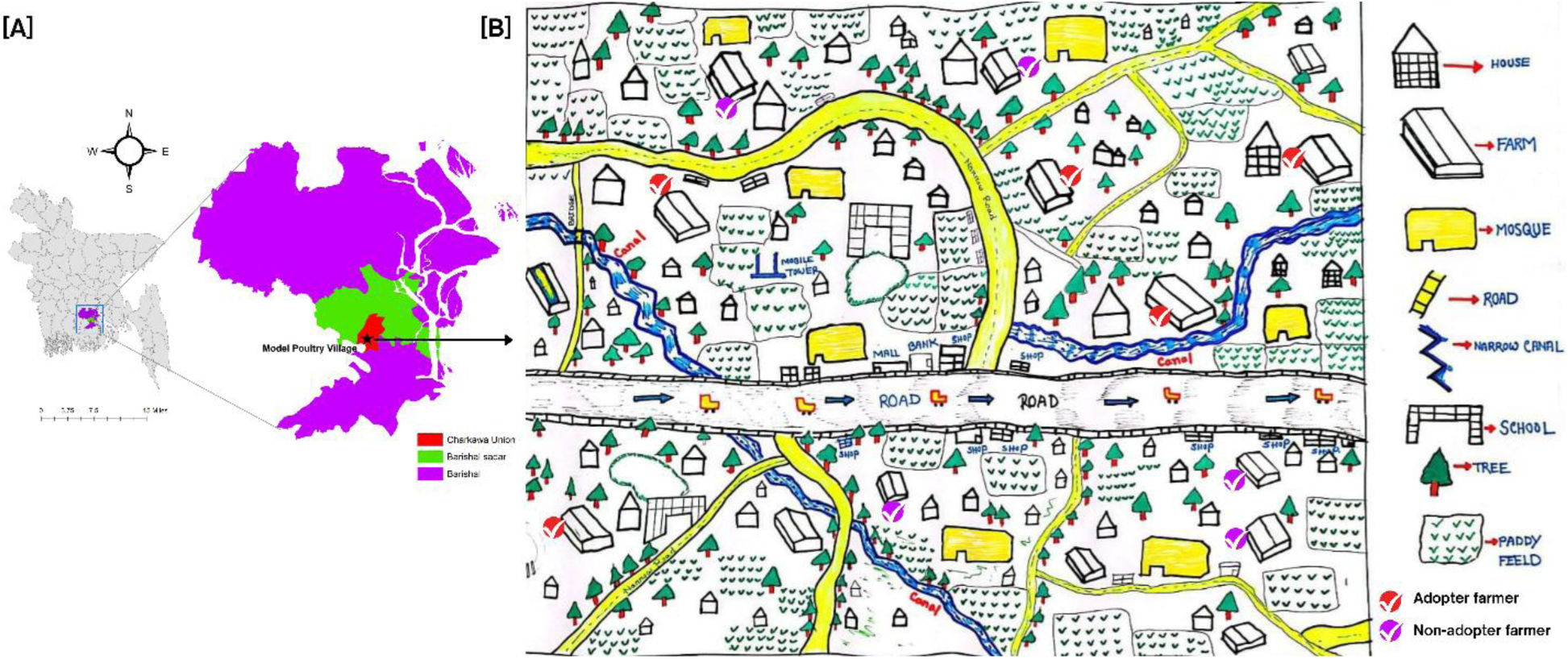
Map of the Model Poultry Village in Barishal Sadar Sub-District, Barishal, Bangladesh. ***Note:*** *(A) Location of the Model Poultry Village in Barishal Sadar sub-district in Barishal. (B) Village map, created by farmers using Participatory Disease Searching (PDS) tools, demonstrates their involvement in mapping and disease monitoring. It shows the village layout with adopter farmers (red arrows) and non-adopter farmers (blue arrows), highlighting key features such as farms, houses, mosques, roads, canals, schools, and paddy fields. Farmers are categorized into adopters and non-adopters for the experimental game study, which will be discussed next in this manuscript*.

### Causal Loop Diagrams (CLDs)

Causal Loop Diagrams (CLDs) were developed through a participatory approach involving 20 broiler farmers, 5 poultry dealers, and 5 veterinary officers from southern Bangladesh. Four discussions were held between September and December 2024 to explore the relationships between antibiotic use, disease vulnerability, and biosecurity practices in poultry farming. Stakeholders were selected through purposive and snowball sampling to ensure relevant experience.

Data collection included semi-structured interviews, focus group discussions, and participatory mapping. In the mapping sessions, stakeholders used large sheets of paper and markers to visually represent key factors such as antibiotic use, disease risk, and biosecurity practices. They drew arrows to show connections between factors, using symbols and notes to clarify the relationships. For instance, they mapped how higher disease risk led to increased antibiotic use and how improved biosecurity could reduce antibiotic dependence.

Stakeholders shared their experiences and perceptions regarding disease outbreaks, antibiotic use, and biosecurity practices. This collaborative process enabled the creation of CLDs, visualizing the interconnected factors influencing disease management in poultry farming. The CLDs were analyzed by identifying key feedback loops and mapping the causal relationships. Positive (+) and negative (−) signs were used to show how changes in one factor affected others within the system.

### Training Approaches and Experimental Game

We conducted six training sessions for 10 farmers at their premises in the Model Poultry Village to present 12 biosecurity and management practices in an engaging and easily understandable manner (**Fig 2, and Fig 3)**. Farmers participated in interactive activities, such as the chili powder game and lipstick demonstration, to learn about germs and the spread of antimicrobial resistance (AMR) through hand contact. In the chili powder game, farmers observed that germs could remain on hands even after washing with water, highlighting the need for disinfection to effectively eliminate germs. The lipstick demonstration showed how AMR could be unknowingly transferred during handshakes. These hands-on exercises emphasized the importance of proper cleaning and biosecurity practices to prevent the entry of germs into poultry farms, thus reducing reliance on antibiotics (**Fig 2**).

**Fig 2.**
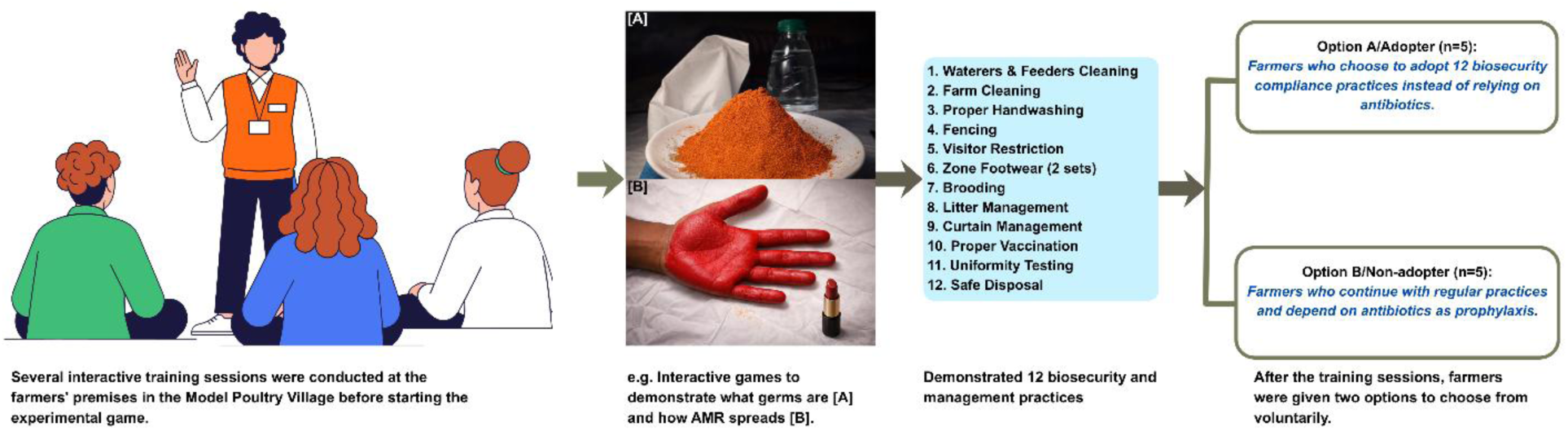
Interactive Training and Experimental Game Setup in the Model Poultry Village.

**Fig 3.**
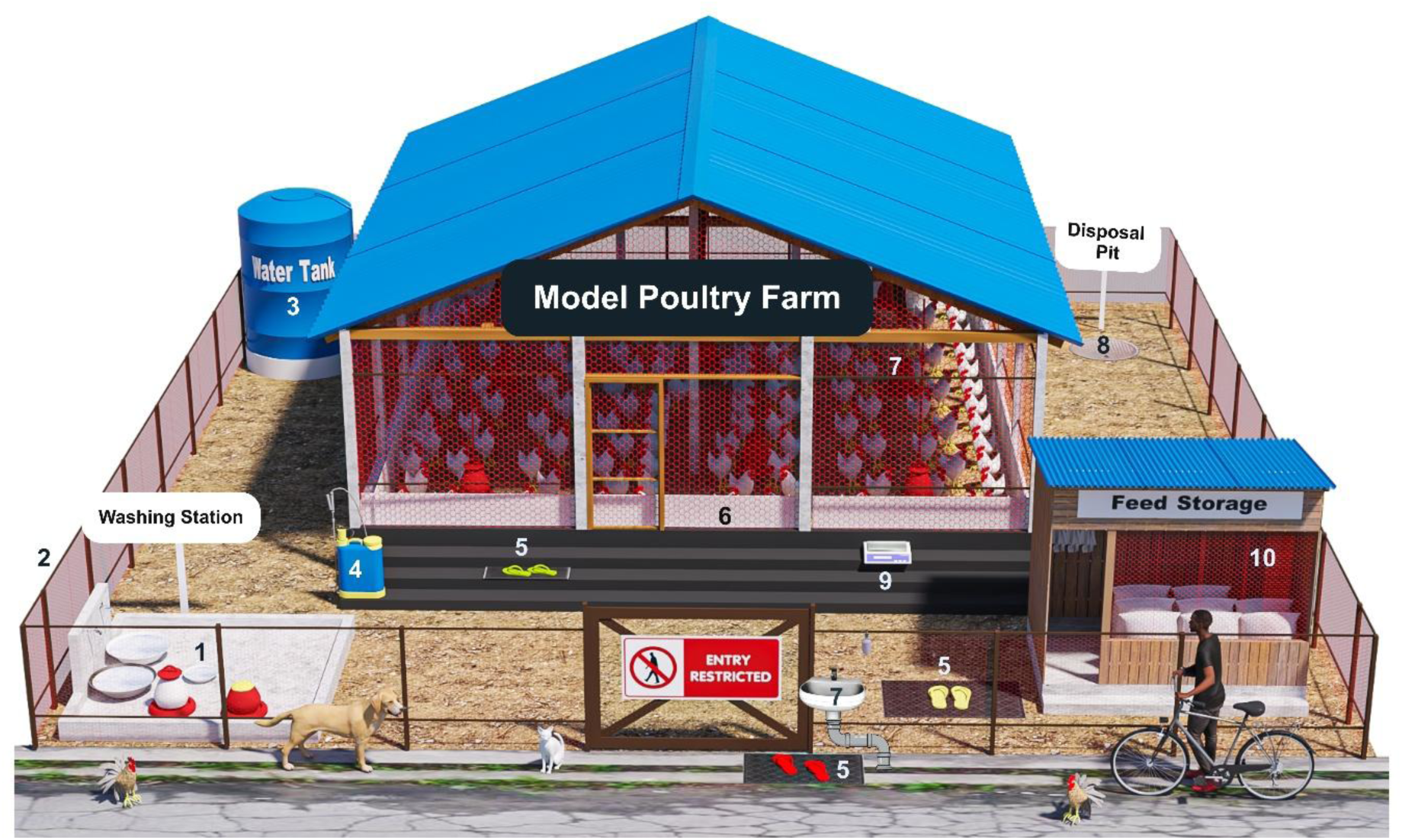
Model broiler poultry house illustrating integrated biosecurity and farm management practices. ***Note:*** *This schematic shows a model broiler poultry farm highlighting key biosecurity and management interventions designed to reduce pathogen entry, improve flock health, and minimize reliance on antibiotics. Numbered labels indicate critical control points within the farm system: (1) dry and wet cleaning of waterers and feeders in a separate washing station; (2) fencing around the farm to prevent entry of visitors, domestic and wild animals, and local and wild birds; (3) ensuring clean and safe water; (4) proper farm cleaning after the end of each batch, maintaining a 15-day gap, and keeping the farm completely locked; (5) zone-specific footwear for cross-contamination prevention (red shoes for regular use outside the farm boundary, yellow shoes for use inside the farm yard, and green shoes for entering the farm); (6) proper curtain management, with curtains hanged at a lower portion; (7) proper brooding management, which is key for optimum production; (8) disposal of dead birds and waste in a designated pit; (9) flock weight and uniformity monitoring at least twice per batch; and (10) maintaining a separate feed storage area*.

To clean the feeder and waterer, farmers were first instructed to remove any dry food residue from the feeder using a brush, then place the residue on white paper and weigh it to calculate feed loss. We mixed disinfectant and water in a bucket, dipped a scrubbing brush into the soapy water, and thoroughly cleaned both the inside and outside of the feeder, ensuring the removal of all biofilm. After rinsing the feeder with clean water, farmers were guided to wipe it with a cloth soaked in germ-killing agents to ensure full coverage. The cleaned feeder was left to dry in the sun. The same process was demonstrated for cleaning the waterer to ensure it was free of water films and germs. Some farmers were surprised by the presence of maggots during dry cleaning, further reinforcing the importance of proper feeder and waterer hygiene. We recommended that farmers use two separate sets of feeders and waterers to maintain better hygiene standards on their farms.

Additionally, we provided hands-on training on effectively spraying germ-killing agents in the poultry shed and instructed farmers to maintain a minimum 15-day gap between batches. Farmers were trained on proper hand-washing techniques, advised to build fences around the farm to keep out surrounding dogs, cats, native chickens, and birds, and encouraged to use at least two pairs of separate shoes for the yellow zone and red zone on the farm. The training covered brooding management, litter management, curtain management, and the proper handling of vaccination and its management. Finally, farmers were instructed on the importance of uniformity testing and the proper disposal of dead birds and waste (**Fig 2, and Fig 3)**.

### Experimental game

After the training sessions, we provided farmers with two options. Option A was for farmers who chose to adopt biosecurity compliance practices instead of relying on antibiotics, while Option B was for farmers who continued with regular practices and depended on antibiotics as prophylaxis. Those who selected Option A were categorized as adopters, while those who chose Option B were categorized as non-adopters. In our experimental game, 5 farmers chose Option A, and 5 farmers chose Option B. Both groups remained stable until Round 4. This division allowed us to assess the impact of adopting biosecurity measures versus continuing with traditional antibiotic use (**Fig 2**).

## Data Variables and Data Collection

Data were collected from 10 farmers before the training and after each of the four training batches to track changes in key variables. To collect baseline data, interviews were conducted using a questionnaire, followed by a visit from a veterinarian. The following variables were recorded after each training batch:

### Biosecurity Practices

Biosecurity practices were assessed on a 0-12 scale by a veterinarian during farm visits.

### Outbreak History

Farmers updated their records with new disease data, which were then verified by a BARA veterinarian during farm visits.

### Mortality Rate

Mortality rate was defined as the proportion of birds that died relative to the total flock size over a given period. Farmers recorded the number of dead birds throughout the study, and mortality rates were calculated at the end of each production batch.

### Antibiotic Use Frequency

Farmers documented the frequency of antibiotic use per flock in their registers, and BARA veterinarians confirmed all prescriptions during visits.

### Feed Conversion Ratio (FCR)

The FCR, calculated as the ratio of feed intake to weight gain, was recorded by farmers during each cycle as part of their routine farm management.

### Profit Per Bird

Profit per bird was calculated by deducting operational costs (DOC, feed, vaccination, medication, electricity, labor and transportation) from the revenue obtained from selling the birds to dealers and consumers.

## Data Analysis

Data collected from all training batches (before and after) were entered into Microsoft Excel for initial data cleaning and organization. Descriptive analysis was performed to evaluate differences between biosecurity adopters and non-adopters, assessing the impact of biosecurity training on key variables such as mortality rate, disease outbreak frequency, antibiotic use frequency, feed conversion ratio (FCR), and profit per bird.

## Software and Tools

ArcGIS 10.8 (Esri, Redlands, USA) was used for visualization and data mapping to track the location of the Model Poultry Village. CANVA and R statistical software (version 4.3.3; R Core Team, 2024) within the RStudio integrated development environment (version 2024.04) were also employed to create visual representations of the data, which contributed to a comprehensive understanding of the impact of biosecurity adoption on poultry production. In addition, Autodesk 3ds Max (Autodesk Inc., San Rafael, USA) with the V-Ray rendering engine (Chaos Group, Prague, Czech Republic) was used to develop a 3D model of the Model Poultry Village to visually illustrate the spatial layout of farms and surrounding infrastructure.

## Ethical Considerations

The study was conducted under the vision of the Department of Livestock Services (DLS), Ministry of Fisheries and Livestock, Government of Bangladesh, with the aim of ensuring the safe, adequate, and high-quality supply of animal products. As part of the annual performance agreement for DLS officials, formal Institutional Review Board (IRB) clearance was not required (Field Disease Investigation Laboratory, Barishal, Department of Livestock Services; Order No. 33.01.0600.415.0001.25.74, Date: 30 May 2025). Written informed consent was obtained from all participants, who were informed of the study’s purpose, procedures, and risks. Participation was voluntary. The study adhered to the Declaration of Helsinki and ethical principles protecting participants’ rights and welfare. It aligned with the One Health approach to improve animal, public, and environmental health. Participant and farm-specific information was anonymized to ensure confidentiality.

## Results

### Causal Loop Diagrams (CLDs)

The Causal Loop Diagram (CLD) illustrates the dynamic relationship between antibiotic use and disease vulnerability in poultry farming. The red reinforcing loop (R1: Antibiotic Dependence) demonstrates how perceived disease risk leads to increased antibiotic use, offering short-term productivity benefits but resulting in long-term vulnerability due to the development of antibiotic resistance. This cycle perpetuates itself, as the increased vulnerability heightens perceived disease risk, driving further antibiotic use. In contrast, the blue balancing loop (B1: Knowledge & Awareness) highlights the role of training and awareness programs. These initiatives encourage farmers to invest in biosecurity measures, reducing reliance on antibiotics and breaking the cycle of dependence. The green loop (B2: Biosecurity & Productivity) shows that biosecurity improvements lower mortality rates, enhance feed conversion ratio (FCR), and increase profitability, which in turn motivates further biosecurity investment. This creates a stabilizing cycle that supports long-term productivity without the need for antibiotics. Finally, the orange loop (B3: Market Incentives) demonstrates how economic incentives, such as demand for antibiotic-free products, encourage farmers to adopt better biosecurity practices, reducing antibiotic dependency. Together, these balancing loops show how awareness, biosecurity, and market incentives can counteract the detrimental effects of antibiotic dependence, fostering a more sustainable and healthier poultry farming system (**Fig 4**).

**Fig 4.**
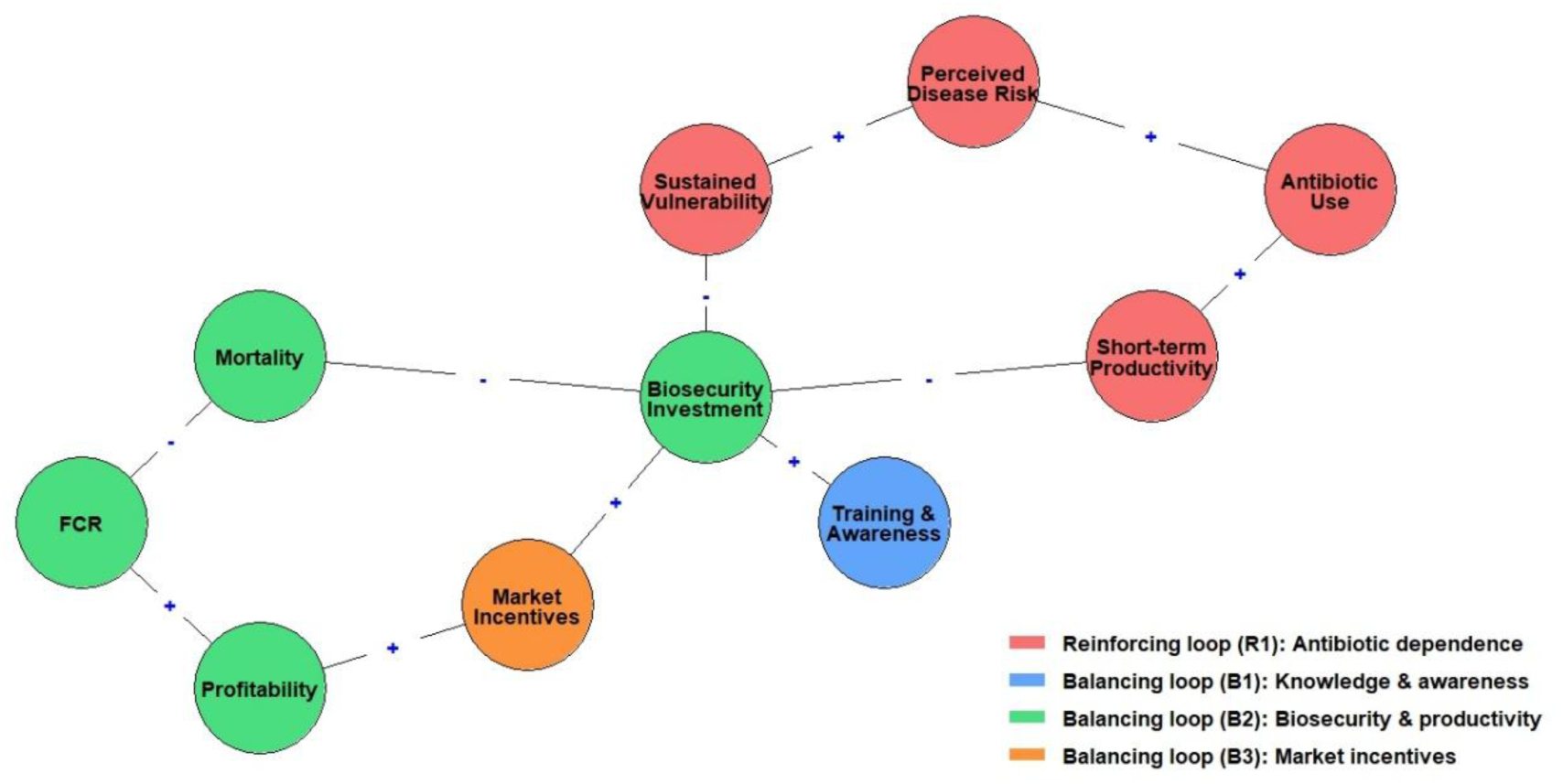
Causal Loop Diagram (CLD) of Antibiotic Use and Biosecurity in Poultry Farming in Bangladesh.

### Demographic and socioeconomic profile of participating farmers

Adopters reared larger flocks (800–2000 birds; mean ≈1000) than non-adopters (600–1000 birds) and were younger (mean 33.6 vs. 38.6 years) with less poultry farming experience (median ≈3 vs. ≈8 years), while non-adopters showed greater variability in experience. Adopters were also more educated, with 80% holding a Hons degree or higher, compared to 40% of non-adopters with SSC or lower, and poultry farming served as the primary income source for 80% of adopters and all non-adopters (**Fig 5**).

**Fig 5.**
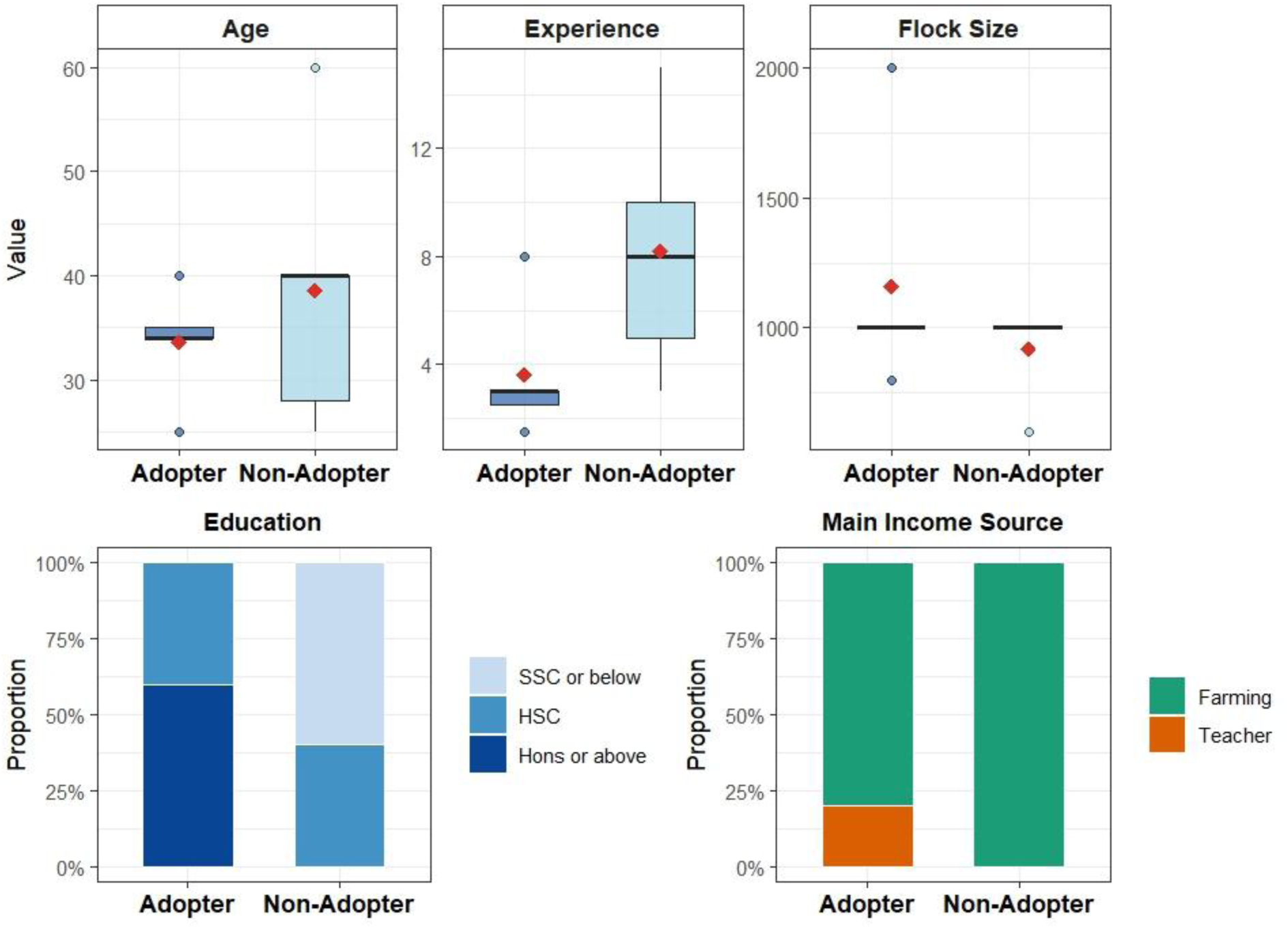
Demographic, educational, and occupational differences between adopters (n=5) and non-adopters (n=5).

### Experimental Games Outcomes

Before training/game (Round 0), both adopters and non-adopters had similar mean biosecurity scores of 0. However, non-adopters outperformed adopters in key variables, including disease outbreaks, mortality, antibiotic use, FCR, and profit per bird. Over the course of the study, adopters consistently improved their biosecurity scores compared to non-adopters, leading to enhanced productivity and profitability. By Round 4, adopters achieved a 125% increase in biosecurity scores (from a mean of 0 to 9, compared to 0 to 4 for non-adopters).

Additionally, adopters experienced a 100% reduction in disease outbreaks (from 2 to 0, compared to 1.4 to 1.0 in non-adopters), a 52.6% reduction in mortality (from 5.6 to 1.8, compared to 4.2 to 3.8 in non-adopters), and a 100% reduction in antibiotic use (from 3.6 to 0, compared to 3 to 2 in non-adopters). FCR for adopters decreased by 13.3% (from 1.8 to 1.3, compared to 1.6 to 1.5 in non-adopters). The profit per bird for adopters increased by 71.72% (from 5.96 to 55.5, compared to 7.5 to 32.3 for non-adopters), reflecting a near-tenfold increase in profitability compared to Round 0 (**Fig 6, and 7)**.

**Fig 6.**
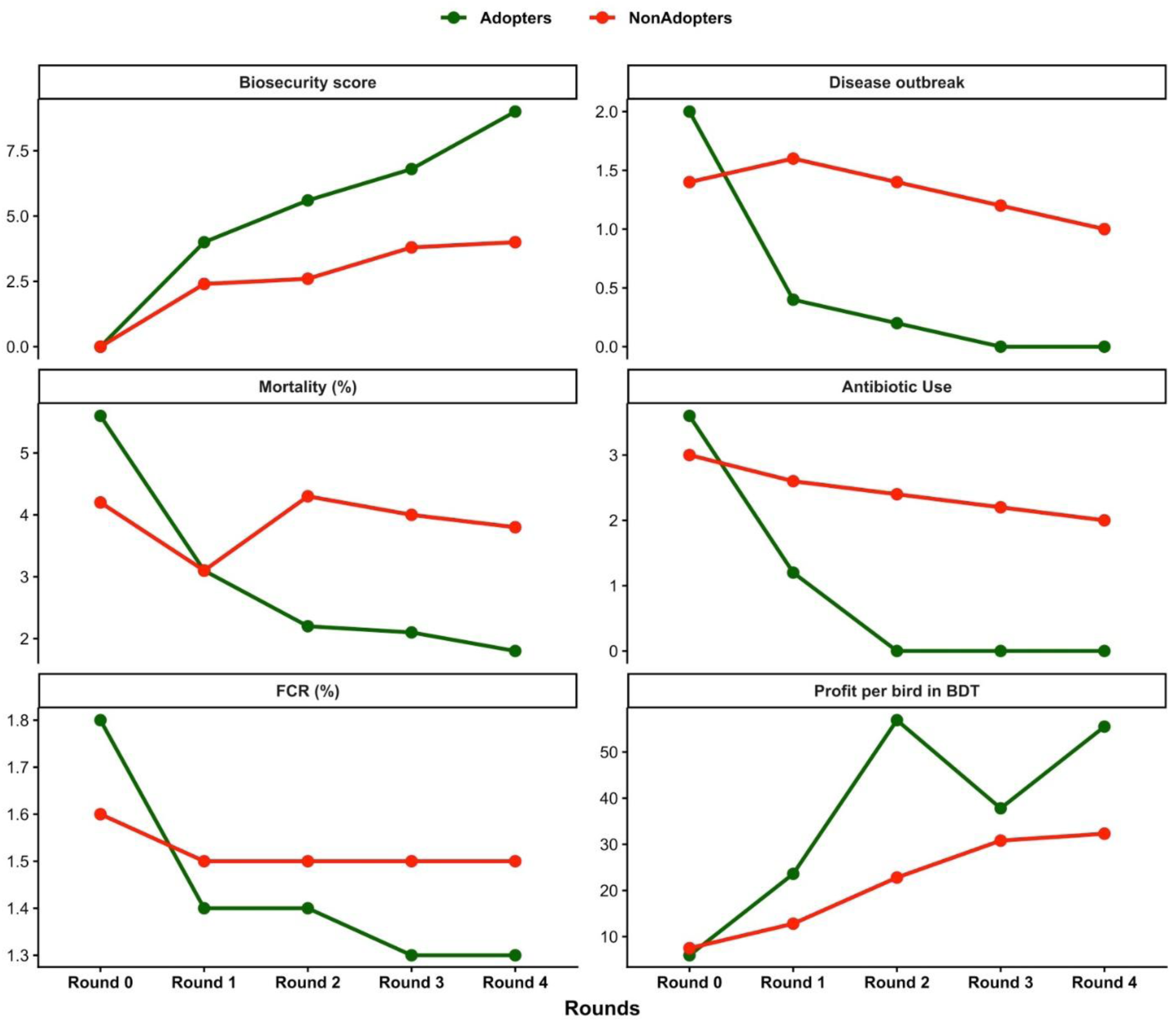
Comparison of biosecurity adoption, antibiotic use, and productivity outcomes for adopters and non-adopters across five rounds. ***Note:*** *Data are mean values across five rounds/poultry rearing batches. Round 0 represents pre-training/game data. Biosecurity is scored on a scale of 1 to 12. Outbreak refers to disease frequency, and Mortality is the percentage of birds lost. Antibiotic Use indicates the quantity of antibiotics used, while FCR (Feed Conversion Ratio) measures feed efficiency as a percentage. Profit is expressed per bird in Bangladeshi Taka (BDT)*.

### Impact of Direct Consumer Sales on Profitability

This figure highlights the significant improvements in farm profitability and health outcomes resulting from biosecurity adoption and direct consumer sales in both the adopter and non-adopter groups. The graph demonstrates that adopters consistently increased the number of birds sold directly to consumers, rising from 31 birds in Round 0 to 60 birds in Round 4, whereas non-adopters remained stable. Dealer prices fluctuated between 128 BDT/kg and 136 BDT/kg across rounds, while consumer prices remained steady at 150 BDT/kg. This price difference allowed farmers to gain extra profit, ranging from 10% to 16% per kg, by selling directly to consumers. The stability in consumer pricing, coupled with increased sales by adopters, reflects a strong strategy for improving profitability over time (**Fig 8**).

**Fig 7.**
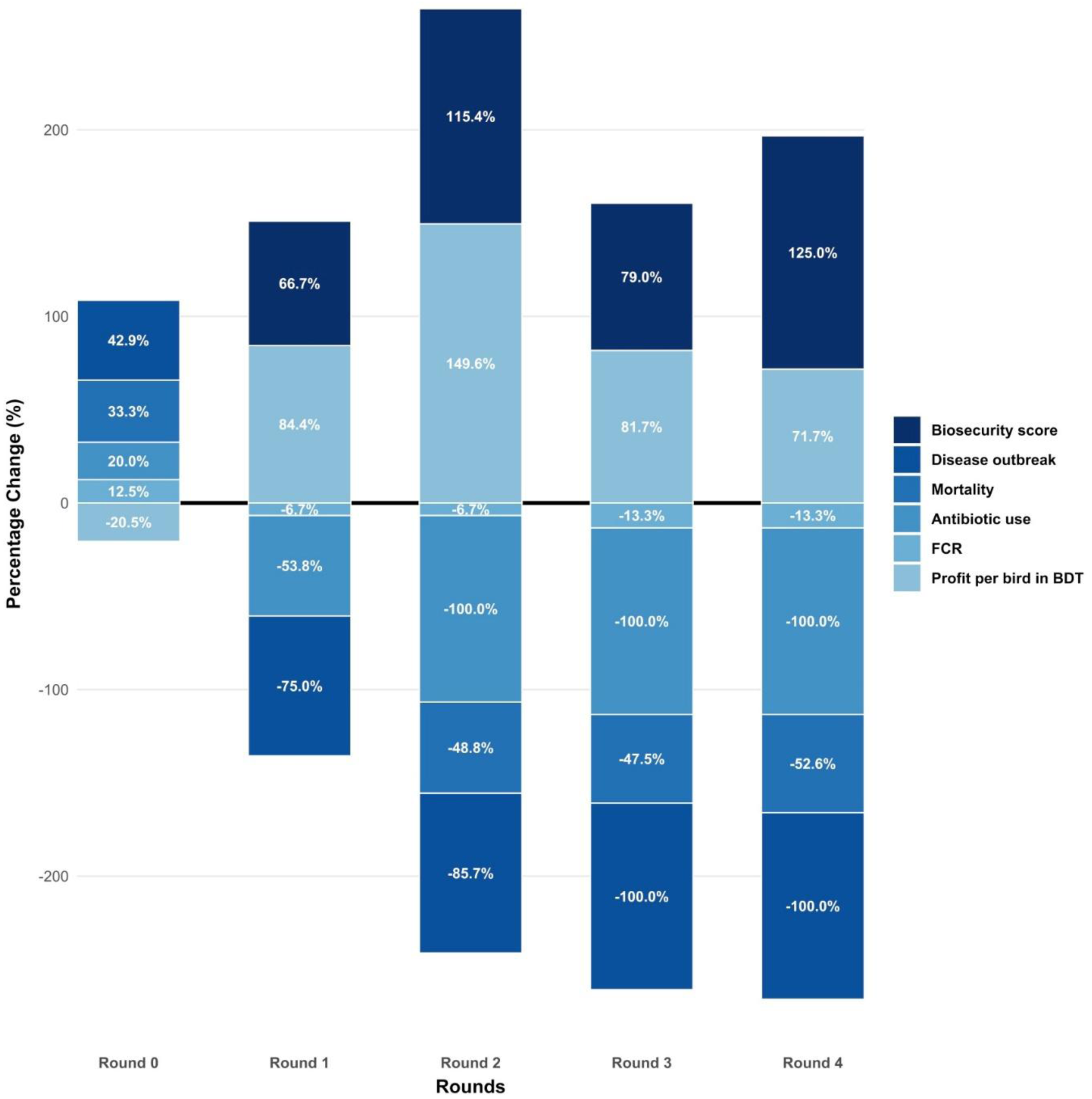
Percentage Changes in Key Variables for Adopters Across Rounds. ***Note:*** *This figure presents the percentage changes in key variables across rounds for adopters compared to non-adopters. Positive values indicate an increase, while negative values denote a decrease in each variable over time. Round 0 represents pre-training/game data. Biosecurity is scored on a scale of 1 to 12. Outbreak refers to disease frequency, and Mortality is the percentage of birds lost. Antibiotic Use indicates the quantity of antibiotics used, while FCR (Feed Conversion Ratio) measures feed efficiency as a percentage. Profit is expressed per bird in Bangladeshi Taka (BDT). The percentage change is calculated by subtracting the value of the non-adopter from the value of the adopter, dividing the result by the value of the non-adopter, and then multiplying by 100*.

**Fig 8.**
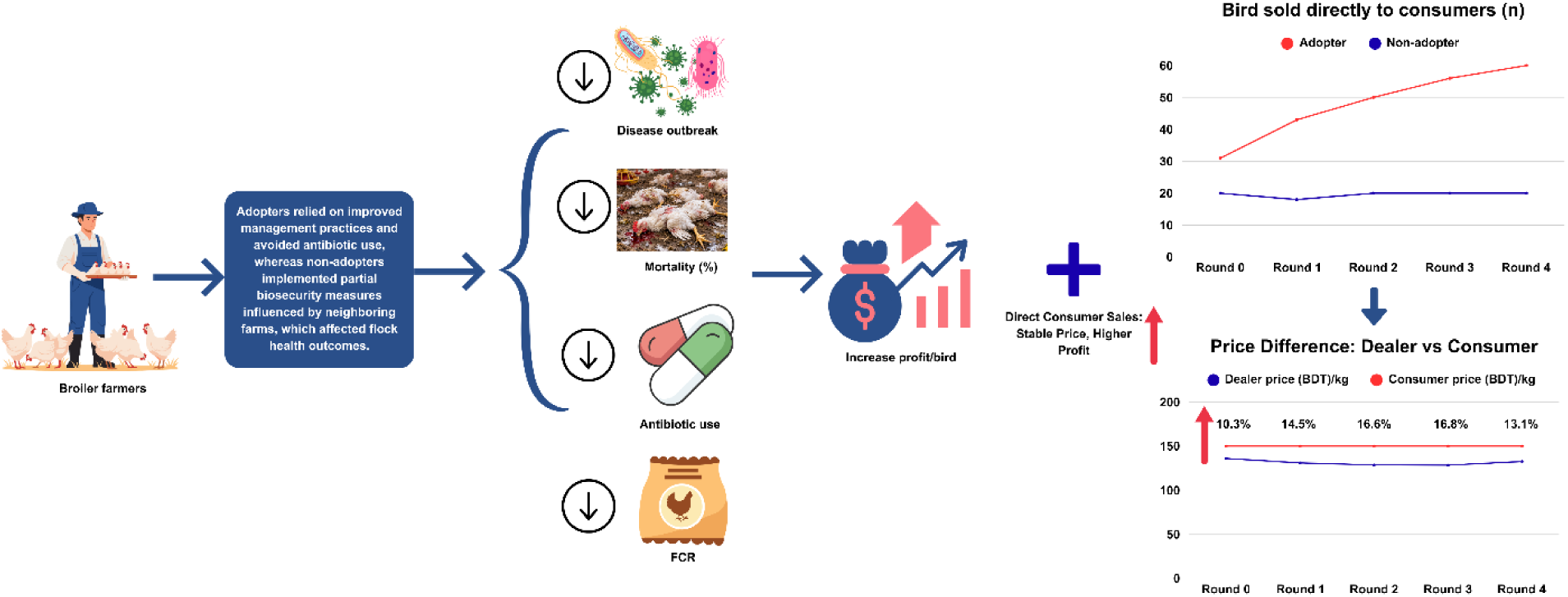
Impact of Biosecurity Practices and Direct Consumer Sales on Farm Profitability and Key Variables.

## Discussion

Antimicrobial resistance (AMR) continues to be a significant global public health threat, particularly in smallholder poultry farming, where antibiotics are often used preventively due to inadequate biosecurity practices [3]. This study highlights how shifting from antibiotic reliance to biosecurity adoption can foster sustainable and resilient poultry systems. While this study provides strong evidence for the effectiveness of biosecurity in reducing antibiotic use, it also raises important questions about the scalability, long-term sustainability, and broader applicability of these practices across different farming contexts.

Causal Loop Diagrams (CLDs) illustrate the dynamics of antibiotic use and disease management. Reinforcing loops show how disease risk perpetuates antibiotic dependence, while balancing loops suggest that biosecurity, education, and market incentives can reduce reliance on antibiotics [33]. In contrast, the balancing loops involving biosecurity practices, farmer education, and market incentives offer a promising pathway to reduce reliance on antibiotics and advance more sustainable practices [34, 35]. Economic pressures, cultural beliefs, and adherence to traditional practices have been shown to limit biosecurity adoption, as reported in studies from India and elsewhere [36, 37]. While the CLDs in this study suggest a promising route for reducing antibiotic use, they do not fully account for the socio-economic barriers that may hinder the scalability of biosecurity practices [38]. Financial constraints, underinvestment in preventive practices, and resistance to changing established routines are obstacles that must be addressed for wider adoption of biosecurity measures in smallholder poultry farming [39, 40].

The demographic and socioeconomic profile of farmers also provides valuable insights into biosecurity adoption. In this study, adopters were younger, more educated, and had less farming experience than non-adopters, consistent with existing literature suggesting that younger, more educated farmers are more likely to adopt innovative practices. A study from Nigeria found that younger farmers, averaging 24 years, showed higher biosecurity adoption rates than older farmers with an average age of 47, despite high awareness of biosecurity principles [41].

Similarly, educated farmers in Pakistan were more inclined to adopt modern agricultural practices [42]. Higher education levels provide farmers with the knowledge to implement biosecurity measures, as seen in Kwara State and Bangladesh, where more educated farmers were more likely to adopt these practices [23, 41]. However, this focus on education highlights a gap in reaching older, less educated farmers, who may perceive biosecurity as costly and time-consuming, contributing to reluctance in adoption [43]. Older farmers also lack access to modern educational resources, further limiting their ability to adopt new technologies [44]. Economic constraints exacerbate this issue, as many farmers prioritize immediate financial returns over long-term investments in biosecurity [39, 40]. Nevertheless, studies suggest that with proper support, older farmers can leverage their experience to implement biosecurity measures when clear benefits, such as cost savings or improved health outcomes, are evident [44]. This indicates that overcoming these barriers through targeted interventions, including educational programs and financial assistance, could enhance biosecurity adoption across various farmer demographics.

In our study, we observed that farmers who were able to achieve higher biosecurity scores showed significant improvements in several key metrics. These farmers experienced fewer disease outbreaks, a reduction in mortality rates with each batch, and a complete elimination of antibiotic use. This reduction in antibiotic use is consistent with findings from previous studies, which demonstrate that stringent biosecurity measures, such as controlled access, disinfection protocols, and waste management, can significantly reduce the incidence of diseases like avian influenza and Newcastle disease [45, 46]. By reducing the chances of disease entry, biosecurity measures lead to fewer outbreaks and lower mortality, a pattern also observed in similar poultry farming contexts [17, 47]. Furthermore, the reduction in antibiotic use improved the Feed Conversion Ratio (FCR), which is often impaired by antibiotics due to the stress they cause in poultry. Stressful conditions reduce feed intake and overall health, leading to less efficient production [48, 49]. As seen in our study, healthier birds had improved FCR due to reduced stress and antibiotic use, which in turn enhanced productivity and reduced pharmaceutical costs. These results are consistent with research showing that biosecurity practices, including reduced antibiotic use, can improve overall flock performance, growth rates, and lower antimicrobial costs [17, 18, 50]. Additionally, the economic benefits of biosecurity were evident, as savings from reduced antibiotic use were reinvested into more sustainable farming practices, leading to increased profit per bird, as reported in studies on the financial advantages of biosecurity [18]. The direct economic and health benefits of biosecurity are further supported by studies showing that biosecured farms often outperform bio-insecured ones in terms of profitability [51].

The results of this study also show that adopters experienced an increase in the sales of their poultry directly to consumers, compared to non-adopters. This increase in sales can likely be attributed to growing consumer awareness of the antibiotic-free (ABF) broilers produced on these farms. As consumers became more aware of the benefits of ABF products, they were more inclined to purchase poultry from these farms. This aligns with findings from previous studies that show a significant preference for ABF products, with over 75% of consumers willing to pay more for pork raised without antibiotics [52]. Furthermore, farmers often sold their poultry to dealers, and the prices they received fluctuated between 128 BDT and 136 BDT. These fluctuations in price, driven by market dynamics, sometimes pushed farmers into a loss-making situation, as seasonal variability and external market forces often cause price instability [53].

However, poultry sold directly to consumers maintained a stable price of 150 BDT across all farms, establishing a more predictable and consistent market. This pricing stability is crucial for farmers, as it reduces income volatility and enhances their financial stability, a benefit already noted in direct-to-consumer (DTC) marketing strategies [54]. If this stable market could be further developed, and antibiotic-free poultry delivered directly to consumers, it would not only ensure better public health outcomes-by reducing antibiotic residues in the food chain [55] but also increase the opportunity for farmers to earn additional benefits [56]. The growing consumer demand for ABF poultry, driven by health consciousness and a desire for safer food options, has been well-documented [57, 58]. Additionally, direct sales foster trust between consumers and producers, as buyers have greater access to information about product origins and safety [59].

This transparency strengthens consumer loyalty and, ultimately, supports more sustainable farming practices. While these findings suggest that biosecurity adoption offers both economic and health benefits, it is important to recognize the broader implications for market development. An interesting aspect of this study is that, before participating in the experimental game, non-adopters were reluctant to adopt biosecurity measures in place of antibiotic dependence, despite being slightly better conditioned than adopters. However, after receiving training, non-adopters began practicing some biosecurity measures, influenced by neighboring adopter farmers, even though they did not change their option in the experimental game. This shift highlights the power of peer influence, as seen in other studies where verbal exchanges and peer interactions significantly contribute to technology adoption. For example, a study found that knowing at least one adopter increased the likelihood of adopting new practices by 26%-32% [60]. Additionally, observing successful practices in neighboring fields boosted adoption rates [61], likely influencing non-adopters in our study to gradually adopt biosecurity measures. These small adaptations resulted in improvements such as reduced disease outbreaks, lower mortality rates, decreased antibiotic use, improved feed efficiency, and increased profit per bird. This aligns with research showing that training and peer influence are crucial for facilitating the adoption of new practices in farming communities [33, 62]. Initially skeptical, non-adopters’ experimentation with biosecurity led to positive feedback and improvements, suggesting that even small changes can yield substantial results. Furthermore, as both adopters and non-adopters saw increased profits, this highlights the potential for the sustainability of biosecurity practices. After the study, all participating farmers agreed that raising broilers without antibiotics was feasible, with some believing it was possible to do so without any medicines. This shift in perception, influenced by adopters, demonstrates the role of social networks in changing beliefs about biosecurity practices. Adopters shared their success stories on social media, encouraging others to explore antibiotic-free poultry farming, reflecting the growing impact of community networks and peer influence in adopting new practices [63]. Trust, built through these networks and reinforced by proximity, plays a key role in adoption [61], and these networks not only foster the adoption of biosecurity but also contribute to its sustainability [64]. These findings align with studies showing the long-term benefits and feasibility of biosecurity measures in poultry farming, highlighting that peer influence and community networks facilitate widespread adoption [65, 66]. While non-adopters initially resisted biosecurity, their gradual adoption, driven by peer influence and experimentation, suggests that community engagement and education are critical for scaling and sustaining biosecurity practices, especially in peripheral farming contexts.

### Limitations

This study has several limitations that should be taken into consideration. First, the sample size was relatively small, comprising only 10 farmers from a single village in Barishal, which may limit the generalizability of the findings to larger or more diverse populations of smallholder poultry farmers. The study’s reliance on self-reported data regarding antibiotic use, mortality rates, and feed conversion ratios introduces potential biases, as farmers may underreport or overreport their practices. Additionally, the study was conducted over a short period, and the experimental game’s duration may not fully capture the long-term effects of biosecurity adoption on farm productivity and sustainability. The absence of a control group limits the ability to compare outcomes between farmers who adopted biosecurity practices and those who did not.

Furthermore, external factors such as regional variations in disease prevalence, market conditions, and farming practices were not controlled for, potentially influencing the results. Lastly, while the study focused on economic and health outcomes, it did not assess the environmental impact of reducing antibiotic use, which could provide a more comprehensive understanding of the broader benefits of biosecurity. Despite these limitations, the study offers valuable insights into the role of training and peer influence in promoting biosecurity adoption. The findings highlight how community-based knowledge transfer can drive behavior change among non-adopters, leading to positive impacts on disease control, productivity, and profitability.

### Future Directions

Future research should aim to extend this study by including a larger and more diverse sample of farmers across different regions of Bangladesh to better assess the broader applicability of these findings. Longitudinal studies are needed to evaluate the long-term sustainability and impacts of biosecurity adoption on farm productivity, health outcomes, and antimicrobial resistance (AMR). Moreover, future studies should incorporate environmental assessments to examine the broader ecological impacts of reduced antibiotic use, including potential reductions in AMR and improvements in soil and water quality. Investigating the role of technology in supporting biosecurity practices, such as mobile applications for monitoring and real-time feedback, would also be a promising direction. Additionally, exploring policy interventions, including financial support or subsidies for biosecurity training, could enhance adoption rates, particularly among resource-constrained farmers. A success story could emerge from this model for developing countries with limited resources, as it shows how reducing antibiotic use at the production level can help mitigate AMR. Finally, addressing barriers to adoption among older, less educated farmers will be essential for scaling biosecurity practices and ensuring equitable benefits across farming communities.

## Conclusion

This study demonstrates that the adoption of biosecurity practices by smallholder poultry farmers can significantly reduce antibiotic use, improve farm productivity, and enhance profitability. The results highlight the critical role of peer influence and community networks in facilitating biosecurity adoption, with non-adopters increasingly practicing biosecurity measures due to the influence of neighboring adopters. Furthermore, consumer demand for antibiotic-free poultry products has provided an additional market opportunity for farmers who adopted biosecurity practices, ensuring better financial stability. However, for biosecurity practices to be widely adopted, it is essential to address socio-economic barriers such as limited access to resources, education, and financial support. By overcoming these barriers and providing continuous education and peer support, biosecurity adoption can become a sustainable and scalable alternative to antibiotic dependence in smallholder poultry farming in developing countries like Bangladesh.

## Acknowledgements

The authors gratefully acknowledge the technical training support provided by the Food and Agriculture Organization (FAO) Emergency Centre for Transboundary Animal Diseases (ECTAD), Bangladesh. Special thanks to the staff of the Field Disease Investigation Laboratory, Barishal, for their valuable technical support. We also thank Dr. Mostafizur Rahman, District Livestock Officer, Barishal; Dr. Kamrun Naher, Ex-Senior National Technical Advisor, ECTAD FAO, Bangladesh; and Dr. Md. Abujar Siddiki, Chief Instructor, Institute of Livestock Science and Technology, Khulna, for their guidance and expertise. Additionally, we appreciate the contributions of the U2C and BARA teams, dealers, veterinarians, and all the participating farmers whose involvement made this study possible.

## References

1. Mestrovic, T., et al., The burden of bacterial antimicrobial resistance in the WHO European region in 2019: a cross-country systematic analysis. The Lancet Public Health, 2022. 7(11): p. e897–e913.

2. Prestinaci, F., P. Pezzotti, and A. Pantosti, Antimicrobial resistance: a global multifaceted phenomenon. Pathogens and global health, 2015. 109(7): p. 309–318.

3. Okeke, I.N., et al., The scope of the antimicrobial resistance challenge. The Lancet, 2024. 403(10442): p. 2426–2438.

4. Murray, C.J., et al., Global burden of bacterial antimicrobial resistance in 2019: a systematic analysis. The lancet, 2022. 399(10325): p. 629–655.

5. Nations, U., Political Declaration of the High-level Meeting on Antimicrobial Resistance. 2024, United Nations General Assembly: New York.

6. Chowdhury, S., et al., Antibiotic usage practices and its drivers in commercial chicken production in Bangladesh. PLoS One, 2022. 17(10): p. e0276158.

7. Raihan, Z., et al., Knowledge, Attitudes and Practices on Antimicrobial Usage and Resistance Among Broiler and Layer Poultry Farmers in Bangladesh: Lessons for Future Improvement. Veterinary Medicine and Science, 2026. 12(1): p. e70776.

8. Masud, A.A., et al., Drivers of antibiotic use in poultry production in Bangladesh: Dependencies and dynamics of a patron-client relationship. Frontiers in Veterinary Science, 2020. 7: p. 78.

9. Faroque, M.O., M.R. Prank, and M. Ahaduzzaman, Effect of biosecurity-based interventions on broiler crude mortality rate at an early stage of production in the small-scale farming system in Bangladesh. Veterinary Medicine and Science, 2023. 9(5): p. 2144–2149.

10. Biswas, P.K., et al., Avian influenza outbreaks in chickens, Bangladesh. Emerging infectious diseases, 2008. 14(12): p. 1909.

11. Bairi, N.S. and S. Beesam, Assessment of Biosecurity Measures in Poultry Farms in and around Warangal District of Telangana State. Indian Journal of Veterinary Public Health| Volume, 2025. 11(1): p. 87.

12. Ziebe, S.D., et al., Impact of biosecurity on production performance and antimicrobial usage in broiler farms in Cameroon. Animals, 2025. 15(12): p. 1771.

13. Islam, M., et al., Poultry Farmer Training in Biosecurity and Production Within an Evaluation Framework in Bangladesh. Veterinary Medicine and Science, 2026. 12(1): p. e70773.

14. Cardona, C.J. and D.R. Kuney, Biosecurity on chicken farms, in Commercial Chicken Meat and Egg Production. 2002, Springer. p. 543–556.

15. Akter, S., et al., Biosecurity practices in commercial chicken farms: Contributing factors for zoonotic pathogen spread. IJID One Health, 2025. 7: p. 100072.

16. Assefa, B., Bio Exclusion and Biocontainment Measures as Effective Poultry Health Management Strategy. J Microbiome Res Health Appl, 2025. 1(1): p. 01–08.

17. Kim, M.B. and Y.J. Lee, Biosecurity Practices for Reducing Antimicrobial Use in Commercial Broiler Farms in Korea. The Journal of Poultry Science, 2025. 62: p. 2025001.

18. Dhaka, P., et al., Can improved farm biosecurity reduce the need for antimicrobials in food animals? A scoping review. Antibiotics, 2023. 12(5): p. 893.

19. Foysal, M., et al., Association between antimicrobial usage and resistance on commercial broiler and layer farms in Bangladesh. Frontiers in Veterinary Science, 2024. 11: p. 1435111.

20. Mendelson, M., et al., Ensuring progress on sustainable access to effective antibiotics at the 2024 UN General Assembly: a target-based approach. The Lancet, 2024. 403(10443): p. 2551–2564.

21. Buchynski, K., S. Jhetam, B.M. Hargis, and K. Schwean-Lardner, Perch use in 11-wk-old turkey hens: impact on performance, health, and behavior. Journal of Applied Poultry Research, 2024. 33(3): p. 100432.

22. Dione, M., et al., *Gendered perceptions of biosecurity and the gender division of labor in pig farming in Uganda.* Journal of Gender, Agriculture and Food Safety, 2020. 5(2): p. 13–26.

23. Alam, M.J., et al., The adoption of biosecurity measures and its influencing factors in Bangladeshi layer farms. Discover Sustainability, 2025. 6(1): p. 29.

24. Sajid, A., N. Kashif, N. Kifayat, and S. Ahmad, Detection of antibiotic residues in poultry meat. Pak. J. Pharm. Sci, 2016. 29(5): p. 1691–1694.

25. Abou-Jaoudeh, C., J. Andary, and R. Abou-Khalil, Antibiotic residues in poultry products and bacterial resistance: A review in developing countries. Journal of Infection and Public Health, 2024. 17(12): p. 102592.

26. Haque, M.H., et al., Sustainable antibiotic-free broiler meat production: Current trends, challenges, and possibilities in a developing country perspective. Biology, 2020. 9(11): p. 411.

27. Haque, C.E. and M. Jakariya, Bengal Delta, char land formation, and riparian hazards: Why is a flexible planning approach needed for deltaic systems? Water, 2023. 15(13): p. 2373.

28. Aktar, S., et al., Geomorphological Responses to Climate Change in the Ganges-Brahmaputra-Meghna Delta: A Multi-Decadal Remote Sensing Analysis. Quaternary Science Advances, 2025: p. 100296.

29. Ali, S.S., et al., Pandemic or environmental socio-economic stressors which have greater impact on food security in the Barishal division of Bangladesh: initial perspectives from agricultural officers and farmers. Sustainability, 2021. 13(10): p. 5457.

30. Ibrahim, M., M. Zaman, A. Mostafizur, and S. Shahidullah, Diversity of crops and land use pattern in Barisal region. Bangladesh Rice J. 21 (2), 57–72. 2018.

31. Nations, F.a.A.O.o.t.U., FAO Impact - Women vaccinators: driving changes in rural Bangladesh. May 14, 2024, FAO: Rome.

32. Thornber, K., et al., Raising awareness of antimicrobial resistance in rural aquaculture practice in Bangladesh through digital communications: a pilot study. Global health action, 2019. 12(sup1): p. 1734735.

33. Guenin, M.-J., M. Studnitz, and S. Molia, Interventions to change antimicrobial use in livestock: A scoping review and an impact pathway analysis of what works, how, for whom and why. Preventive veterinary medicine, 2023. 220: p. 106025.

34. Vougat Ngom, R., et al., Resistance to medically important antimicrobials in broiler and layer farms in Cameroon and its relation with biosecurity and antimicrobial use. Frontiers in Microbiology, 2025. 15: p. 1517159.

35. Arjmand, A., et al., Assessing the impact of biosecurity compliance on farmworker and livestock health within a one health modeling framework. One Health, 2025. 20: p. 101023.

36. Mehmedi, B., et al., Economic Perspectives on Farm Biosecurity: Stakeholder Challenges and Livestock Species Considerations. Agriculture, 2025. 15(21): p. 2288.

37. Panda, P., et al., Awareness and adoption of farm biosecurity practices in commercial dairy, pig and poultry farms of Uttar Pradesh (India). Tropical Animal Health and Production, 2024. 56(6): p. 203.

38. Saccavini, O., et al. Towards collective biosecurity: a participatory approach in French communities of poultry farmers. in First conference on Animal Biosecurity WABA. 2025.

39. Buckel, A., et al., Understanding the factors influencing biosecurity adoption on smallholder poultry farms in Ghana: a qualitative analysis using the COM-B model and theoretical domains framework. Frontiers in Veterinary Science, 2024. 11: p. 1324233.

40. Naumovska, S., et al., Biosecurity measures for prevention and control of Salmonella in poultry farms. 2025.

41. Tsado, J., et al., Determinants of adoption of biosecurity principles by poultry farmers in Kwara State, Nigeria. Ethiopian Journal of Environmental Studies & Management, 2017. 10(4).

42. Basil, M., U. Muzammal, and U. Ahmad, Poultry farmers’ perceptions and practices regarding the use of growth promoters in commercial broiler production in Punjab, Pakistan. Sci Soc Insights, 2025. 1(2): p. 66–74.

43. Damiaans, B., S. Sarrazin, E. Heremans, and J. Dewulf, Perception, motivators and obstacles of biosecurity in cattle production. Vlaams Diergeneeskundig Tijdschrift, 2018. 87(3).

44. Lestari, V., D. Rahardja, and S. Sirajuddin. Barriers to adopt biosecurity at smallholder farmers. in IOP Conference Series: Earth and Environmental Science. 2022. IOP Publishing.

45. Islam, Z., et al., Assessment of biosecurity measures in commercial poultry farms of Rajshahi district in Bangladesh. Preventive Veterinary Medicine, 2023. 219: p. 106027.

46. Islam, M., M. Meher, A. Harun, and M. Haider, The Common respiratory diseases of Poultry in Bangladesh: present status and future directions. Veterinary Sciences: Research and Reviews, 2022. 8(1): p. 52–64.

47. Mallioris, P., et al., Biosecurity and antimicrobial use in broiler farms across nine European countries: toward identifying farm-specific options for reducing antimicrobial usage. Epidemiology & Infection, 2023. 151: p. e13.

48. Lara, L.J. and M.H. Rostagno, Impact of heat stress on poultry production. Animals, 2013. 3(2): p. 356–369.

49. Wasti, S., N. Sah, and B. Mishra, Impact of heat stress on poultry health and performances, and potential mitigation strategies. Animals, 2020. 10(8): p. 1266.

50. Jaramillo, S., *Biosecurity: A holistic One-Health concept for enhancing the health, welfare, and productivity of commercial poultry-A point of view.* Proceeding of The First International Avian Influenza Summit, University of Arkansas-October, 2023: p. 16–17.

51. Fasina, F.O., et al., The cost–benefit of biosecurity measures on infectious diseases in the Egyptian household poultry. Preventive veterinary medicine, 2012. 103(2-3): p. 178–191.

52. Kozera-Kowalska, M. and J. Uglis, Consumer attitudes and purchase intentions for antibiotic-free pork in Poland: An empirical study. 2024.

53. Das, S., et al., Predicting and mitigating agricultural price volatility using climate scenarios and risk models. arXiv preprint arXiv:2503.24324, 2025.

54. Key, N., Direct-to-Consumer Marketing and the Survival and Growth of Beginning Farms. Journal of Agricultural and Resource Economics, 2024. 49(1): p. 81–101.

55. Izah, S.C., et al., Public health risks associated with antibiotic residues in poultry food products. Journal of Agriculture and Food Research, 2025. 21: p. 101815.

56. Muaz, K., et al., Antibiotic residues in chicken meat: Global prevalence, threats, and decontamination strategies: A review. Journal of food protection, 2018. 81(4): p. 619–627.

57. Zhou, Y., A. Zhang, R.D. van Klinken, and J. Wang, Understanding consumers’ purchase intention towards meat produced without preventive antibiotic use. Foods, 2024. 13(23): p. 3779.

58. Mohammadi, H., S. Saghaian, and F. Boccia, Antibiotic-free poultry meat consumption and its determinants. Foods, 2023. 12(9): p. 1776.

59. Moustier, P. and T.T.L. Nguyen, Direct sales suit producers and consumers’ interests in Vietnam, in From Community to Consumption: New and Classical Themes in Rural Sociological Research. 2010, Emerald Group Publishing Limited. p. 185–197.

60. Massfeller, A. and H. Storm, Field observation and verbal exchange as different peer effects in farmers’ technology adoption decisions. Agricultural Economics, 2024. 55(5): p. 739–757.

61. Bandiera, O. and I. Rasul, Social networks and technology adoption in northern Mozambique. The economic journal, 2006. 116(514): p. 869–902.

62. Dione, M.M., et al., Impact of participatory training of smallholder pig farmers on knowledge, attitudes and practices regarding biosecurity for the control of African swine fever in Uganda. Transboundary and emerging diseases, 2020. 67(6): p. 2482–2493.

63. Yadav, J., et al., Role of social media in technology adoption for sustainable agriculture practices: Evidence from twitter analytics. Communications of the Association for Information Systems, 2023. 52(1): p. 833–851.

64. Kolady, D., W. Zhang, T. Wang, and J. Ulrich-Schad, Spatially mediated peer effects in the adoption of conservation agriculture practices. Journal of Agricultural and Applied Economics, 2021. 53(1): p. 1–20.

65. Achoja, F., Cost and Return Analysis of Biosecurity Management in Poultry Farms in Rivers State, Nigeria. Black Sea Journal of Agriculture, 2020. 3(1): p. 43–48.

66. Otieno, W.A., R.A. Nyikal, S.G. Mbogoh, and E.J. Rao, Adoption of farm biosecurity practices among smallholder poultry farmers in Kenya–an application of latent class analysis with a multinomial logistic regression. Preventive Veterinary Medicine, 2023. 217: p. 105967.

